# A Novel Approach to Combat *Pseudomonas aeruginosa*: Repurposing Pharmaceuticals for Inhibition of Phospholipase A

**DOI:** 10.1101/2025.03.23.644859

**Authors:** Matea Modric, Rocco Gentile, Lena Schröder, Raphael Moll, Ifey Alio, Wolfgang R. Streit, Karl-Erich Jaeger, Holger Gohlke, Filip Kovacic

**Affiliations:** Institute of Molecular Enzyme Technology, Heinrich Heine University Düsseldorf, Forschungszentrum Jülich GmbH, D-52425 Jülich, Germany; Institute for Pharmaceutical and Medicinal Chemistry, Heinrich Heine University Düsseldorf, 40225 Düsseldorf, Germany; Department of Microbiology and Biotechnology, University of Hamburg, Ohnhorststr. 18, 22609 Hamburg, Germany; Institute of Bio- and Geosciences (IBG-4: Bioinformatics), Forschungszentrum Jülich GmbH, 52425 Jülich, Germany; Department of Surgery, Massachusetts General Hospital, Boston, Massachusetts, USA and Department of Microbiology, Harvard Medical School, Boston, Massachusetts, USA

## Abstract

Phospholipases A (PLAs) play critical roles in cellular physiology, making human PLAs established drug targets. On the other hand, the potential of bacterial PLAs as targets for antimicrobial drug development remains underexplored. In this study, we curated a library of 23 approved and investigational pharmaceuticals, some of which inhibit human PLA-like enzymes, through a combination of ligand structure-based searches and textual mining in literature and compound databases. Experimental screening identified that compounds GW4869, darapladib, and rilapladib significantly inhibit *Pseudomonas aeruginosa* growth by more than 50 %. While these compounds did not reduce biofilm formation, GW4869 increased the proportion of dead cells in established biofilms, suggesting its role in compromising biofilm cell viability. Biochemical assays revealed that all three compounds inhibited the enzymatic activity of PlaF, a PLA virulence factor of *P. aeruginosa*, by decreasing the affinity of a model substrate. Molecular dynamics simulations and binding free energy analyses indicate that GW4869 binds to the substrate-binding and product-release tunnels of PlaF, suggesting GW4869 as a non-covalent competitive inhibitor. Notably, the mutant strain *P. aeruginosa* Δ*plaF* proved to be GW4869 resistant and did not display differential growth upon GW4869 treatment, further indicating PlaF as the primary GW4869 target. Furthermore, GW4869 and rilapladib significantly enhanced the efficacy of the last-resort antibiotic imipenem in combination treatments. These findings highlight the potential of GW4869, darapladib, and rilapladib to act as repurposed inhibitors of PlaF or PLA-dependent mechanisms in *P. aeruginosa* and underscore the promise of combination therapies against intracellular PLAs to combat antimicrobial resistance.

## 1. Introduction

*Pseudomonas aeruginosa* is a nosocomial Gram-negative pathogen that poses a significant threat to public health, particularly among immunocompromised patients (Reynolds and Kollef 2021, Jesudason 2024). Its multidrug resistance has led the World Health Organization (WHO) to classify it as a priority pathogen requiring the urgent development of effective antibiotics and alternative treatment strategies (Murray, Ikuta et al. 2022; Melchiorri, Rocke et al. 2024). The overuse and misuse of antibiotics, inadequate waste management, and rapid environmental transmission have further exacerbated antibiotic resistance in *P. aeruginosa* and other ESKAPE pathogens (Qin, Xiao et al. 2022, Elfadadny, Ragab et al. 2024). To address the global resistance crisis that hinders the effectiveness of existing antibiotics against *P. aeruginosa* (Miethke, Pieroni et al. 2021), recent advances have been made in the development of innovative antivirulence approaches against this human pathogen (Veesenmeyer, Hauser et al. 2009; Qin, Xiao et al. 2022; Jiang, Zheng et al. 2024).

These antivirulence compounds target key virulence determinants, reducing pathogenicity or enhancing susceptibility to antibiotics, thereby improving infection treatment outcomes (Kudoh, Wiener-Kronish et al. 1994; Kurahashi, Kajikawa et al. 1999; Sadikot, Blackwell et al. 2005; Kipnis, Sawa et al. 2006; Jiang, Zheng et al. 2024). Among them, the most promising approaches include inhibition of quorum sensing (Müh, Schuster et al. 2006; Singh, Almpani et al. 2022), surface-associated adhesins (Kao, Churchill et al. 2007), and iron acquisition pathways (Choi, Britigan et al. 2019) that collectively contribute to biofilm formation (Zheng, Raudonis et al. 2019), inhibition of secretory systems (Neely, Holder et al. 2005) and direct neutralization of extracellular toxins such as phospholipase A (PLA) (Lee, Pukatzki et al. 2007). However, monotherapy with these compounds has shown limited success in eradicating infections (Zheng, Raudonis et al. 2019, Wang, Lu et al. 2024). Combination therapies, such as quorum sensing inhibitors and tobramycin or the iron chelator deferoxamine and tobramycin (Moreau-Marquis, O’Toole et al. 2009), have demonstrated enhanced efficacy in eliminating *P. aeruginosa* biofilms in infected hosts. These findings underscore the potential of adjuvant therapies targeting virulence factors to become integral components of clinical practice (Veesenmeyer, Hauser et al. 2009; Chatterjee, Anju et al. 2016).

Among the virulence factors of *P. aeruginosa*, secreted PLA play an important role in host cell invasion, membrane disruption, and immune evasion by releasing fatty acids from host phospholipids (Flores-Díaz, Monturiol-Gross et al. 2016). Notably, pseudolipasin A, a selective inhibitor of the ExoU phospholipase, has shown protective effects *in vitro* on hamsters, amoebae, and yeast cells against ExoU-mediated toxicity (Lee, Pukatzki et al. 2007). Beyond secreted phospholipases, emerging evidence reveals the significance of intracellular bacterial phospholipases for virulence (Bleffert, Granzin et al. 2022, Caliskan, Poschmann et al. 2023) and antibiotic resistance (Kerrinnes, Young et al. 2015). Their function is related to the adjustment of the bacterial membrane phospholipid composition to adapt to environmental stimuli. For example, *Brucella melitensis* PLA BveA increases the resistance to polymyxin B through the hydrolysis of membrane phosphatidylethanolamine (Kerrinnes, Young et al. 2015). Similarly, the intracellular PLA PlaF of *P. aeruginosa* regulates membrane phospholipid composition, thereby influencing iron uptake, motility, biofilm formation, and signaling pathways (Caliskan, Poschmann et al. 2023). PlaF-deficient strains exhibit reduced virulence in mouse macrophage and *Galleria mellonella* infection models, underscoring its therapeutic potential (Bleffert, Granzin et al. 2022; Kovacic and Batra-Safferling, 2025).

The crystal structure of PlaF reveals a transmembrane helix anchoring the catalytic domain, which contains a serine-hydrolase catalytic triad, to the cytoplasmic membrane (Bleffert, Granzin et al. 2022). This configuration facilitates the interaction of PlaF with membrane phospholipid substrates and enables substrate extraction via one of three distinct active-site tunnels (Ahmad, Strunk et al. 2021; Gentile, Modric et al. 2024). The crystallized complex of PlaF with the fatty acid product and molecular dynamics simulations identified putative product egress tunnels (Ahmad, Strunk et al. 2021, Gentile, Modric et al. 2024). Detailed structural and mechanistic insights furthermore emphasize the drugability of PlaF.

In light of the challenges posed by traditional drug discovery, this study explores the repurposing of FDA-approved or clinically tested compounds as potential inhibitors of PlaF. Some of the selected compounds have been described to target human serine hydrolase family PLAs (Bachovchin and Cravatt 2012). From a library of 23 identified compounds, we experimentally tested their effects on *P. aeruginosa* PA01 growth, biofilm formation and dispersal. Growth-inhibiting compounds were further assessed for PlaF inhibition using functional enzyme assays and a Δ*plaF* mutant strain. Enzyme kinetics and molecular dynamics simulations indicate the mode of action of the PlaF-targeting compound GW4869. Experiments to mimic combination therapy demonstrated synergism between GW4869 and the last-resort antibiotic imipenem supported the therapeutic potential of PlaF as an antivirulence target. This study underscores the inhibition of intracellular PLAs as a novel and previously underexplored strategy for developing effective treatments against *P. aeruginosa*.

## 2. Material and methods

### 2.1. Protein expression and purification

Expression and purification of PlaF were performed as described previously (Bleffert, Granzin et al. 2019). *P. aeruginosa* PA01 cells were transformed (Choi, Kumar et al. 2006) with plasmids pBBR-*plaF_H6_* or pBBR-*plaF_H6_*-F229W (Kovacic, Bleffert et al. 2016) and grown overnight at 37 °C in lysogeny broth (LB) medium supplemented with tetracycline (100 µg/mL). These cultures were used to inoculate an expression culture in LB medium to an initial optical density at 600 nm (OD_600nm_) of 0.05 at 600 nm. The cultures were grown at 37 °C until OD_600nm_ ∼ 2 and afterward harvested by centrifugation at 6750 × *g* and 4 °C for 15 min. The total membrane fraction was solubilized with Triton X-100, and proteins were purified using Ni-NTA agarose (Qiagen, Hilden, Germany) and buffer supplemented with 0.25 mM *n*-dodecyl-β-D-maltoside (DDM). For biochemical analysis, proteins were transferred to Tris-HCl buffer (100 mM, pH 8) supplemented with the respective detergent. The purity of PlaF was analyzed by sodium dodecyl sulfate-polyacrylamide gel electrophoresis (SDS-PAGE) under denaturation conditions on a 12 % (w/v) gel (Laemmli 1970).

### 2.2. Selection and preparation of compounds for screening library

The compound library was assembled based on the 2D structural similarity to six approved pharmaceuticals identified through a literature search as indicated in Table 1. The DrugBank database was queried with these six selected compounds, and structures with more than 50 % 2D structural similarity to the initial compounds were included in the library. Additionally, the Selleckchem database was searched using the keyword “phospholipase” and the phospholipase inhibitors were included to the library. The stock solutions of selected compounds were prepared using eiter dimethyl sulphoxide (DMSO), methanol or water as indicated in Table S1.

**Table 1.**
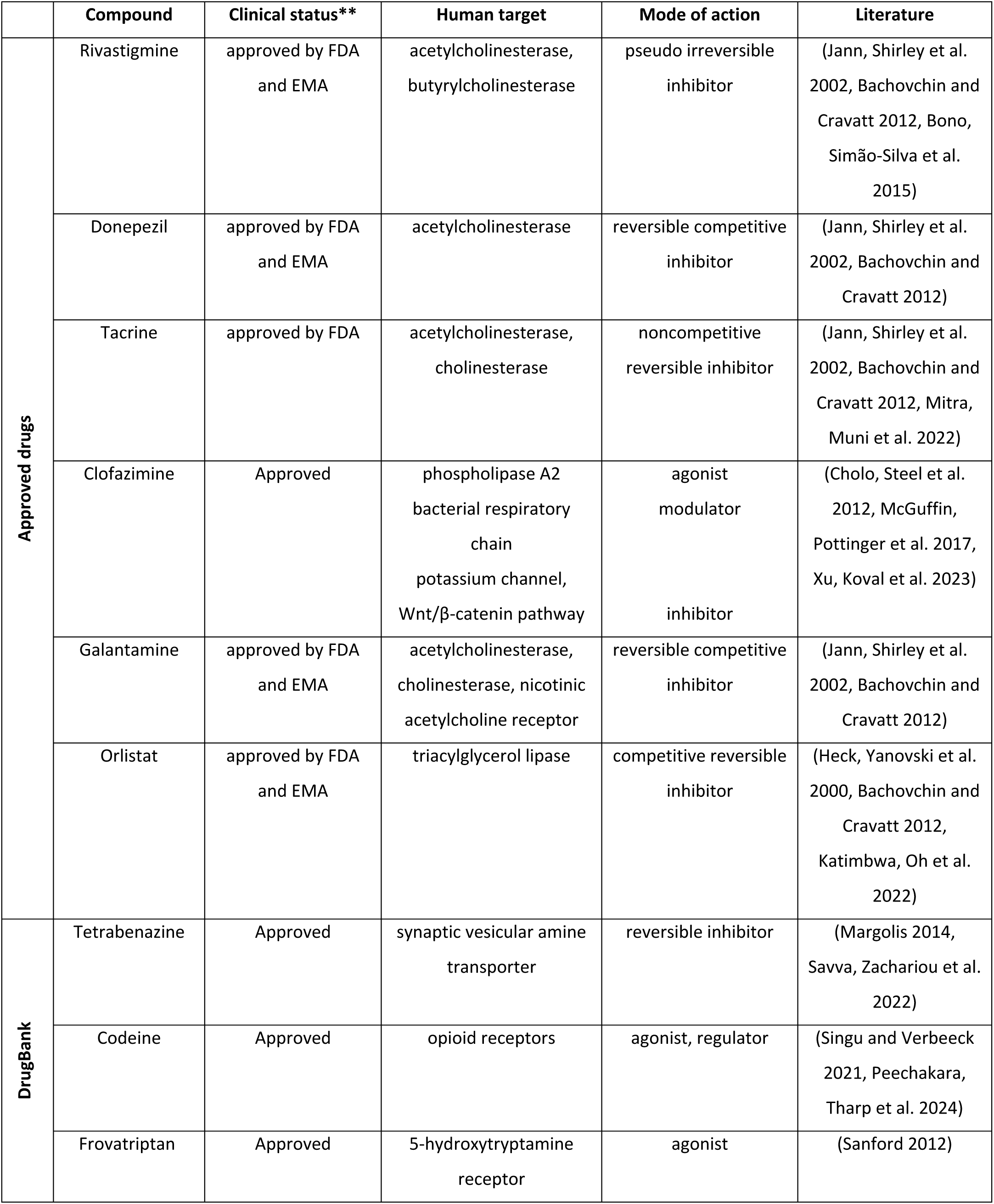
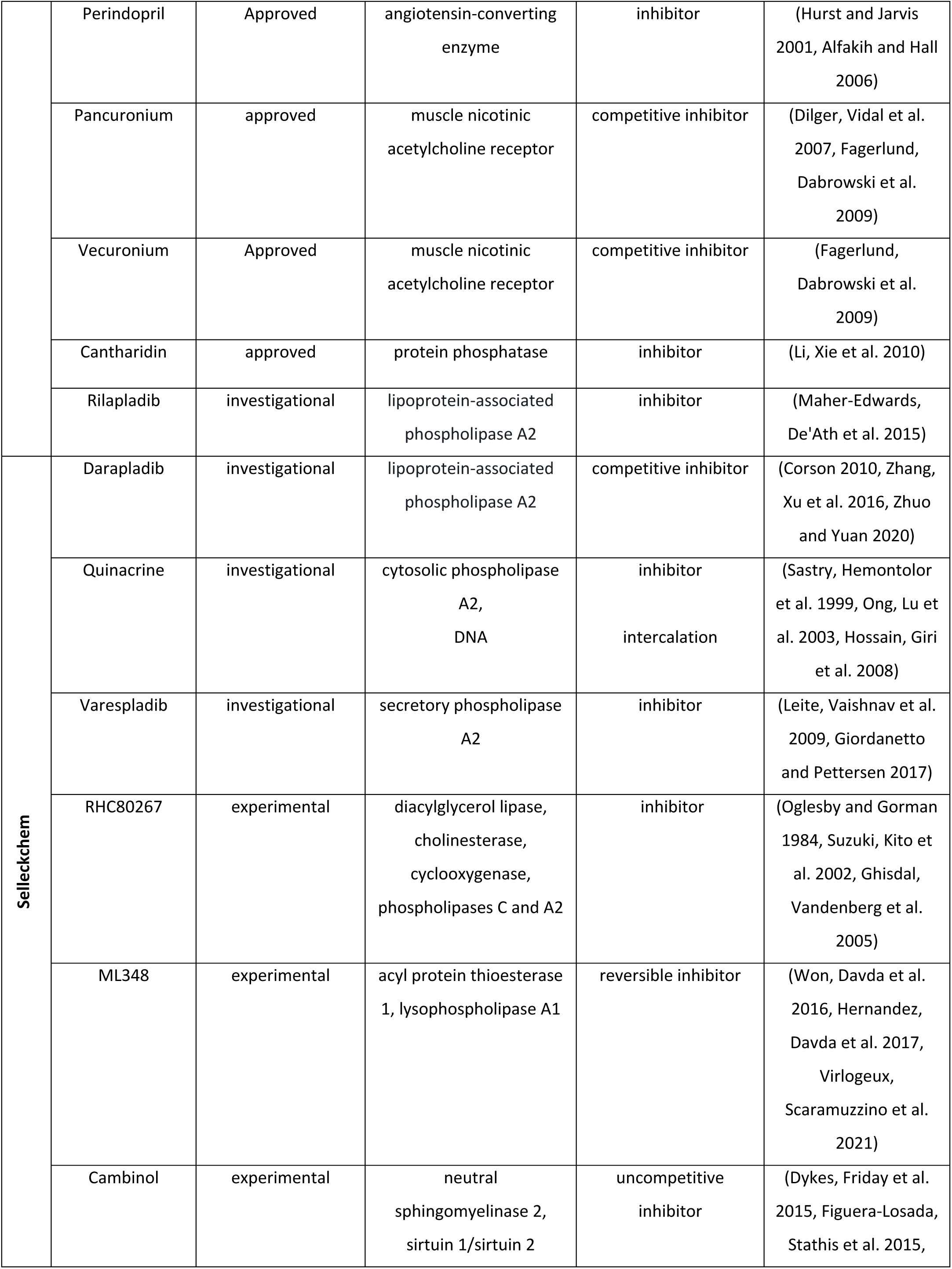
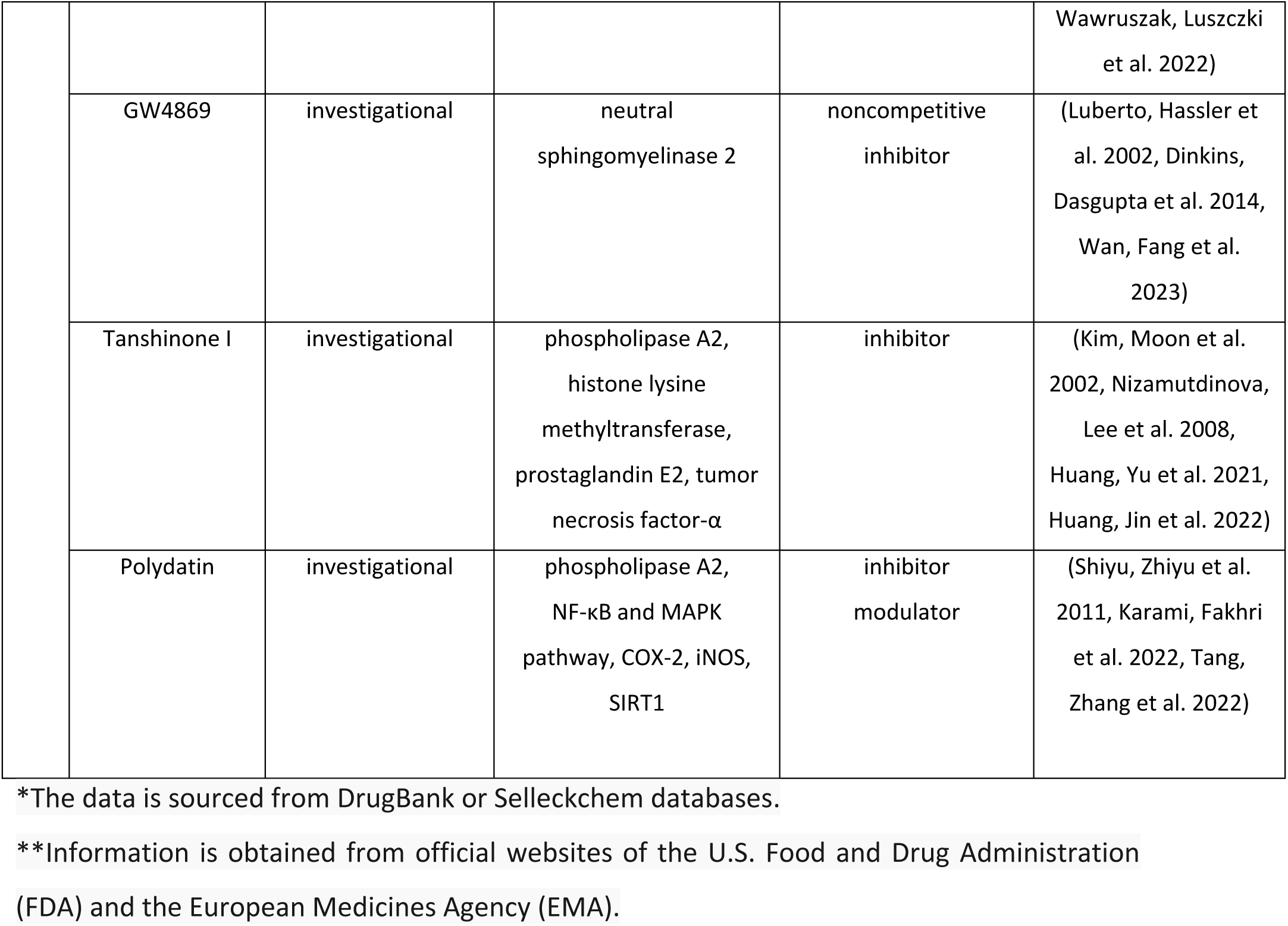
Library of pharmaceuticals*.

### 2.3. Growth curves

The growth of bacterial strains *P. aeruginosa* PA01 wildtype an isogenic mutant Δ*plaF* (Bleffert, Granzin et al. 2022) was monitored by measuring the OD_600nm_ for 10 h in a 96-well plate shaken with agitation of 1000 rpm. First, the OD_600nm_ of overnight cultures was diluted with lysogeny broth medium to 0.05. When the bacterial cultures reached an OD_600nm_ of approximately 0.4, 1.5 µL of inhibitor stock solutions (Table S1), 1.5 µL of antibiotic, or both were added in a total volume of 150 µL. The antibiotics were prepared in water at the following final concentrations: 0.5 mg/L (0.70 µM) for gentamicin, 2 mg/L (3.71 µM) for piperacillin, 1 mg/L (0.71 µM) for colistin, and 2 mg/L (6.30 µM) for imipenem. Cultures treated with respective solvents were used as no-compound controls.

### 2.4. Enzyme activity assay, inhibition, and enzyme kinetic studies

The esterase activity of the PlaF protein was determined spectrophotometrically with *p*-nitrophenyl butyrate (*p*-NPB) as a substrate, as described previously (Jaeger and Kovacic 2014), using a 96-well microplate. 5 µL of purified protein (stock concentration 30 µg/mL) was mixed with 2 µL of potential inhibitor stock solution at concentrations of: 1.73 mM for GW4869, 3.34 mM for codeine, and 10 mM for all the remaining compounds and 93 µL of freshly prepared 1 mM *p*-NPB solution was added. Esterase activity was monitored by increase of absorbance at 420 nm for upto 2 h at 37 °C. Kinetic parameters were determined by measuring activity with different substrate concentrations (0.05, 0.1, 0.2, 0.3, 0.5, 1, 1.3, 1.5 mM) (Bleffert, Granzin et al. 2022). Half-maximal inhibitory concentration (IC_50_) was determined by measuring enzyme activity with different compound concentrations and with 1 mM *p*-NPB substrate and was calculated from linear plots (Jayachandra, Gowda et al. 2023).

### 2.5. Biofilm assay

Microtiter dish biofilm assay was performed according to a modified protocol (Reisner, Krogfelt et al. 2006; O’Toole 2011). The respective number of single colonies of *P. aeruginosa* was inoculated in LB media and incubated for 24 h at 37 °C with shaking. Overnight cultures were used to inoculate 150 µL of bacterial cultures of OD_600_ = 0.05 in LB medium in a plastic 96-well microtiter plate. In the case of compound-treated biofilm, 2 µL of the compound or respective solvent were added to a total volume of 150 µL. The inoculated microtiter plate was covered with air-permeable sealing film and incubated for 24 h at 37 °C. Afterwards, the OD_600nm_ was measured, medium was removed, each well was briefly washed two times with 200 µL LB media at 37 °C. Biofilms were stained by adding 200 µL 0.1 % (w/v) crystal violet solution, followed by 20 minutes incubation at room temperature. The crystal violet solution was discarded and the plate was washed three times with 200 µL water, followed by adding 200 µL 30 % (v/v) acetic acid. The plate was incubated for 20 minutes at room temperature, and absorbance was measured at 585 nm using a plate reader (SpectraMax iD3, Molecular devices GmbH, München, Germany).

### 2.6. Colony forming unit count

Biofilm was grown as described above. After 24 h incubation, the supernatant was discarded, the biofilm was suspended in 200 µL LB medium, diluted 100,000 fold, and 100 µL of cell suspension was plated on LB medium agar plates. Plates were incubated overnight at 37 °C.

### 2.7. Quantification of extracellular DNA in biofilm

Extracellular DNA was quantified according to a modified protocol (Leggate, Allain et al. 2006). Biofilm was grown in a 96-well plate as described above. After washing two times with 200 µL LB medium, 100 µL of TE buffer (10 mM Tris-HCl, 1 mM EDTA, pH 8) was added, followed by adding 100 µL of SYBR Green I (ThermoFisher Scientific, Germany). SYBR Green I was prepared freshly by diluting it 1:1250 in TE buffer. The plate was incubated for 10 minutes in the dark at room temperature, and the fluorescence was measured using a fluorescence plate reader (SpectraMax iD3, Molecular devices GmbH, München, Germany) at 485/518 nm excitation/emission filter set up.

### 2.8. Confocal laser scanning microscopy of biofilm

Confocal laser scanning microscopy (CLSM) of biofilm was performed as described previously (Alio, Gudzuhn et al. 2020). Briefly, bacterial biofilms were prepared under static conditions in 8-well chamber µ-slide (ibidi GmbH, Gräfelfing, Germany) and treated with the compounds listed in Table S1. After 24 h incubation, the samples were incubated with live/dead staining solution (LIVE/DEAD BacLight^TM^ Bacterial Viability Kit, Thermo Fisher Scientific, Waltham, USA) to assess the viability of the bacterial cells. The biofilms were visualized by “Confocal laser scanning microscope 800” (Axio observer.Z1/7, Carl Zeiss AG, Oberkochen, Germany) at settings listed in Table S2.

### 2.9 Preparation of starting structures for unbiased molecular dynamics simulations

The crystal structure of the PlaF dimer (PDB ID 6I8W) is available in the Protein Data Bank (Berman 2000). The last five residues of the C-terminus of each monomer missing in the structure were added using MODELLER (Sali and Blundell 1993), and all small molecule ligands were removed. The dimer was oriented in the membrane using the PPM server (Lomize, Pogozheva et al. 2012). From that, the monomeric configuration of PlaF chain A was generated by removing chain B from the dimer orientation. Chain A was oriented again using the PPM server resulting in the tilted configuration t-PlaF.

The structure of GW4869 was prepared starting from its canonical SMILES using the fixpka option in OpenEye (<u>OEDocking 4.3.0.3;</u> https://www.eyesopen.com). The most favorable conformer was generated with Omega 4.1.1.1 (OMEGA 4.1.1.1; http://www.eyesopen.com) using the flag –maxconfs = 1. The charges of GW4869 were calculated following the RESP approach (Schauperl, Nerenberg et al. 2020) using Gaussian 16 to compute electrostatic potentials at the HF/6-31G* level (Frisch 2016). GW4869 docking to t-PlaF was performed using Autodock-3.0.5 (Morris, Goodsell et al. 1998) with the in-house developed scoring function DrugScore (Dittrich, Schmidt et al. 2019). The selected configuration represents the most populated cluster obtained.

The bound t-PlaF configuration was embedded into a DOPE:DOPG = 3:1 membrane (Murzyn, Róg et al. 2005) and solvated using PACKMOL-Memgen (Martínez, Andrade et al. 2009, Schott-Verdugo and Gohlke 2019). The membrane composition resembles that of the native inner membrane of Gram-negative bacteria (Murzyn, Róg et al. 2005) and was already used for simulating PlaF in a membrane bilayer environment (Ahmad, Strunk et al. 2021; Bleffert, Granzin et al. 2022, Gentile, Modric et al. 2024). A distance of at least 15 Å between the protein or membrane and the solvent box boundaries was kept. To obtain a neutral system, counter ions were added that replaced solvent molecules (KCl 0.15 M) resulting in the systems containing ∼140,000 atoms.

### 2.10 Unbiased molecular dynamics simulations of selected inhibitors bound to t-PlaF

The GPU particle mesh Ewald implementation from the AMBER23 suite of molecular simulation programs (Case, Aktulga et al. 2023) with the ff14SB (Maier, Martinez et al. 2015), Lipid21 (Dickson, Walker et al. 2022), and GAFF2 (He, Man et al. 2020) force fields for the protein, membrane lipids, and ligands, respectively, were used; water molecules and ions were parametrized using the TIP3P model (Zhao, Zhao et al. 2019) and the Li and Merz 12-6 ions parameters (Li, Song et al. 2015, Sengupta, Li et al. 2021). For each t-Plaf-inhibitor complex, five independent replicas of 1 μs length were performed. Covalent bonds to hydrogens were constrained with the SHAKE algorithm (Kräutler, Van Gunsteren et al. 2001) in all simulations, allowing the use of a time step of 2 fs. Details of the thermalization of the simulation systems are given below. All unbiased molecular dynamics (MD) simulations showed structurally invariant protein structures and membrane phases evidenced by electron density calculations (Figure S1). The RMSD of GW4869, after removal of the global motions of t-PlaF, shows structurally invariant binding configurations across five different replicas (Figure S2).

### 2.11 Relaxation, thermalization, and production runs

An initial minimization step was performed with the CPU code of pmemd (Le Grand, Götz et al. 2013). Each minimization was organized in four steps of 1000 cycles each, for a total of 4000 cycles of minimization. Afterward, each minimized system was thermalized in one stage from 0 to 300 K over 25 ps using the NVT ensemble and the Langevin thermostat (Quigley and Probert 2004), and the density was adapted to 1.0 g cm^−3^ over 4975 ps using the NPT ensemble with a semi-isotropic Berendsen barostat (Berendsen, Postma et al. 1984), with the pressure set to 1 bar. The thermalization and equilibration were performed with the GPU code of pmemd (Le Grand, Götz et al. 2013).

For each replica, 1 μs of production run using the GPU code of pmemd was performed in the NPT ensemble at a temperature of 300 K using the Langevin thermostat (Quigley and Probert 2004) and a collision frequency of 1 ps^−1^. To avoid noticeable distortions in the simulation box size, semi-isotropic pressure scaling using the Berendsen barostat (Berendsen, Postma et al. 1984) and a pressure relaxation time of 1 ps was employed by coupling the box size changes along the membrane plane (Lin, Pan et al. 2017).

### 2.12 MM-PBSA calculations of GW4869 binding to t-PlaF

To pinpoint the most likely binding epitopes, we generated 3D density grids to map the location of GW4869 across multiple replicas. We considered stably bound conformations if a frame of the trajectory has a ligand RMSD < 1.5 Å to the previous frame (see Supplementary results). These binding configurations were clustered using the hierarchical agglomerative (bottom-up) algorithm implemented in cpptraj (Roe and Cheatham 2013), using the minimum distance *ε* between the clusters as the cluster criterion. Starting from *ε* = 2.0 Å, we gradually increased *ε* in 0.5 Å intervals until the population of the largest cluster remained unchanged (at *ε* = 4.0 Å). We calculated the 3D density maps of GW4869 considering all atoms using the grid function available in cpptraj (Roe and Cheatham 2013) with a grid spacing of 1.5 Å. We applied a contour level of 1*σ* (one standard deviation above the mean value). Subsequently, we conducted MM-PBSA (Molecular Mechanics-Poisson Boltzmann Surface Area) calculations with MMPBSA.py (Miller, McGee et al. 2012) and Normal Mode Analysis computations to determine the binding effective energy and the binding entropy of GW4869 to t-PlaF, respectively. These calculations were performed across the different replicas, and the results per replica were averaged. Dielectric constants of 1.0 for the protein, 80.0 for the solvent, and 15.0 for the membrane were used, similar to our previous work (Gentile, Modric et al. 2024). A heterogeneous dielectric model was used to represent the membrane; the implicit membrane model using spline fitting (memopt = 3) was employed for the binding effective energy calculations (Greene, Qi et al. 2019). This allowed us to compute Δ*G*_solv_ and Δ*G*_gas+solv_ (eq. 1-2) (Gohlke and Case 2004). The normal mode analysis calculation allowed us to calculate the loss of flexibility upon binding -*T*Δ*S*_total_ considering the translational, rotational, and vibrational terms (eq. 3) (Gohlke and Case 2004).

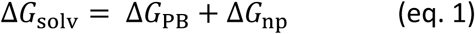

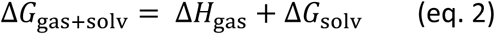

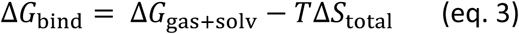

To compare the computed with experimentally determined binding affinities, we converted Δ*G*_bind_ into the standard free energy of binding 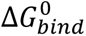 according to eq. 4, as done previously (Gohlke, Kiel et al. 2003; Frieg, Gremer et al. 2020). This takes into account that translational entropy depends on solute concentration (McQuarrie 1976; Janin 1996), leading to the concentration dependence of chemical equilibria that do not conserve the number of molecules (such as binding reactions) (Gilson, Given et al. 1997; Luo and Sharp 2002).

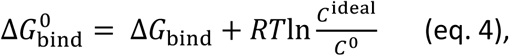

where *R* is the universal gas constant (*R* = 0.001987 kcal K^−1^ mol^−1^), *T* = 298.15 K, *C*^0^ is the standard concentration of 1 mol l^−1^, and *C*^ideal^ the ligand concentration of 0.041 mol l^−1^, derived from the general gas equation at a pressure of 101,325 Pa and a temperature of 298.15 K (Gohlke, Kiel et al. 2003). 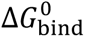 is directly related to the computed dissociation constant 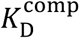 according to eq. 5.

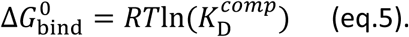

*K*_D_ values derived from computations according to eq. 2-5 are denoted as 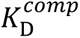. The total standard error of the mean of the computations, denoted as *SEM*_total_, is estimated following the principles of Gaussian error propagation according to eq. 6.

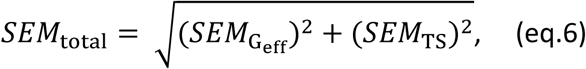

where *SEM*_Geff_ and *SEM*_TS_ are the SEMs from MM-PBSA and NMA computations, respectively. The results from binding free energy calculations are reported as 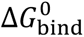 ± *SEM*_total_. The computations were converged, as evidenced by the comparison between the first and second halves of the trajectories (Figure S3).

## 3. Results

### 3.1. Identifying pharmaceuticals with potential inhibitory effects on bacterial phospholipases A

This study aimed to identify approved drugs and pharmaceuticals from (pre)clinical studies that have potential inhibitory activity against the bacterial phospholipases PlaF to evaluate their suitability for repurposing as antimicrobials. In the following, we will jointly refer to these compounds as pharmaceuticals. According to the literature, the six FDA-approved drugs, rivastigmine (Jann, Shirley et al. 2002; Bachovchin and Cravatt 2012), donepezil (Jann, Shirley et al. 2002; Bachovchin and Cravatt 2012), galantamine (Jann, Shirley et al. 2002; Bachovchin and Cravatt 2012), tacrine (Jann, Shirley et al. 2002; Bachovchin and Cravatt 2012), orlistat (Heck, Yanovski et al. 2000; Bachovchin and Cravatt 2012), and clofazimine (Cholo, Steel et al. 2012; McGuffin, Pottinger et al. 2017), target human PLAs or esterases/lipases, enzymes known for their side PLA activity due to the shared serine hydrolase catalytic mechanism (Kovacic, Bleffert et al. 2016; Bleffert, Granzin et al. 2022). These pharmaceuticals were used as templates to identify similar compounds among more than 500,000 investigational, clinically tested, or approved drugs in the DrugBank database (Wishart 2006; Wishart, Knox et al. 2008). This ligand-based search identified eight additional compounds showing 2D structural similarity with the initial six compounds, with similarity scores ranging from 54 to 72 %. Furthermore, a targeted keyword search in the Selleckchem database, which includes over 120,000 bioactive small-molecules, yielded nine additional compounds being investigated as human phospholipase, lysophospholipase, or lipase inhibitors (Figure 1).

**Figure 1.**
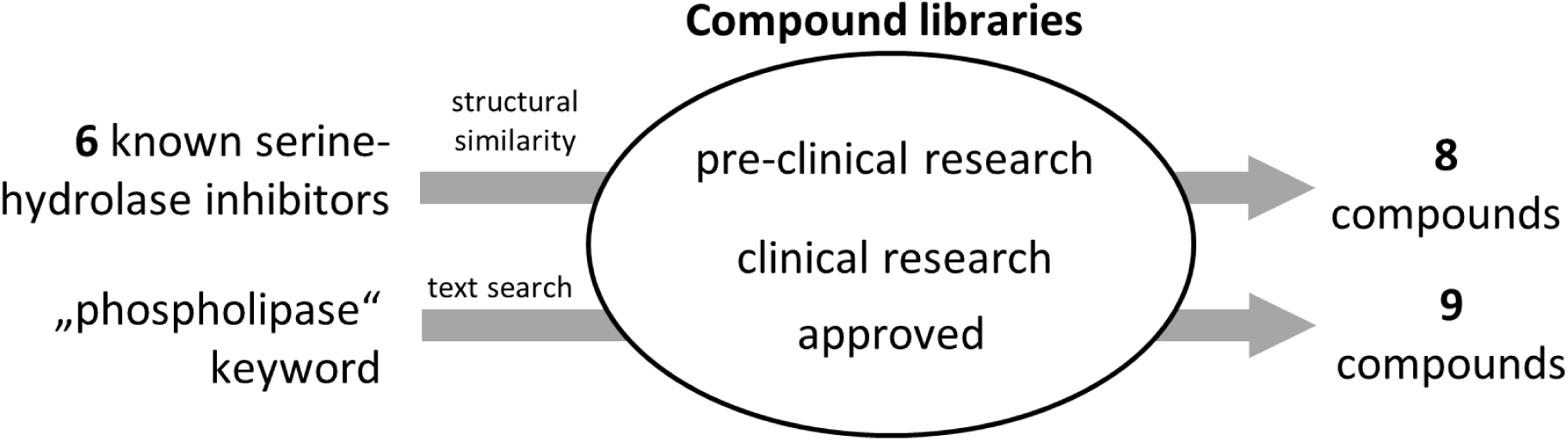
Selection of potential phospholipase PlaF inhibitors. A library of 23 pharmaceuticals to be tested as potential PlaF inhibitors (Table 1) was generated by screening compounds in preclinical and clinical research phases and approved drugs, from compound libraries of DrugBank and Selleckchem databases. The selection criteria involved a combination of structural similarity- and text-based searches.

Most of the resulting 23 pharmaceuticals, among which thirteen have regulatory approval, seven are in clinical research phases, and three are in preclinical studies, target esterases, lipases, and phospholipases. For the cases of tetrabenazine, codeine, frovatriptan, perindopril, pancuronium, vecuronium, and cantharidin, no related target has been described, despite an apparent molecular similarity to the template compounds or associated keywords. The advantage of this library is that the majority of the compounds have met safety and efficacy standards for use in humans (Table 1). In conclusion, this library of diverse potential phospholipase inhibiting compounds provided a foundation for testing their repurposing as antimicrobial agents.

### 3.2 Effect of potential PlaF inhibitors on the growth and biofilm formation of *P. aeruginosa*

We investigated the effects of the 23 pharmaceutical compounds from the library on the planktonic growth of *P. aeruginosa* in a rich LB medium. In a screening experiment designed to identify the most potent pharmaceuticals, the compounds were added to exponentially growing bacterial cultures (OD_600nm_ ∼ 0.5) in plastic microtiter plates (MTP), and optical density was measured after incubation at 37 °C with agitation. The results demonstrated that GW4869, and rilapladib significantly inhibited bacterial growth compared to the respective solvent-treated controls after 5 h incubation, while the effect by darapladib is significant after 7 h (Figure 2A). Validation growth inhibition experiments under these screening conditions revealed that the effects of all three compounds were evident as early as 2-4 h post-treatment (Figure 2B). Over 6–7 h, treated cultures exhibited a slower growth rate compared to the controls. After 6–7 hours, darapladib and rilapladib reduced *P. aeruginosa* growth significantly by approximately 35 %, while GW4869 exhibited a more pronounced effect, reducing growth by 57 % (Figure 2B).

**Figure 2.**
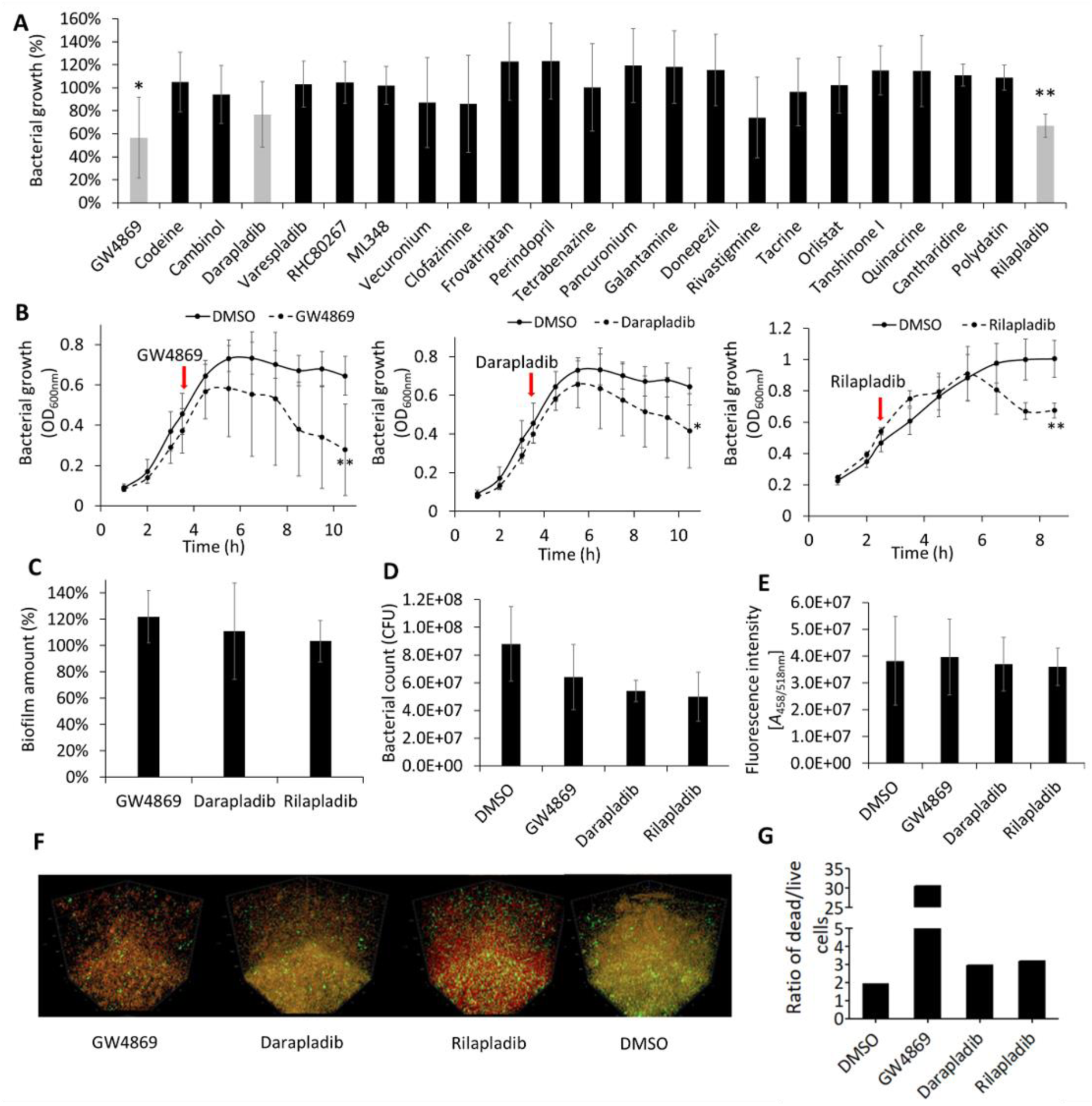
Potential PLA inhibitors impact the growth and viability of *P. aeruginosa* PA01. **A)** Effect of 23 pharmaceuticals on the planktonic growth of *P. aeruginosa* after 5 h incubation with compounds. Relative growth values represent optical densities (OD_600nm_) of compound-treated cultures compared to the OD_600nm_ of respective solvent-treated cultures set to 100 %. Light grey bars represent pharmaceuticals that significantly impacted the growth of *P. aeruginosa* after 5 h or more (see panel B). Black bars represent compounds that showed no significant effect on the *P. aeruginosa* growth at any time point. Results are shown as the mean ± S.D. of 6 biological replicates (n = 6) from two independent experiments. *t*-test of normally distributed values, \**p* < 0.05, \*\**p* < 0.01. **B)** Growth curves of *P. aeruginosa* treated with GW4869 (17.3 µM), darapladib (100 µM), and rilapladib (100 µM). Bacterial cultures were grown in LB medium at 37 °C with shaking at 1000 rpm in plastic MTP. Red arrows indicate beginning of treatment. Results are the mean ± S.D. of three independent experiments (n = 3). *t*-test of normally distributed values, \**p* < 0.05, \*\**p* < 0.01. **C)** Biofilm amount of *P. aeruginosa* treated for 24 h with GW4869 (17.3 µM), darapladib (100 µM), and rilapladib (100 µM) was quantified by the crystal violet assay. The results are shown relative to the solvent-treated cultures, which were set to 100 %. The results are the mean ± S.D. of > 22 biological replicates from four independent experiments. *t*-test was calculated compared to the untreated samples. **D)** Viable bacterial count determined as colony forming units (CFUs) within biofilm treated for 24 h with GW4869 (17.3 µM), darapladib (100 µM), and rilapladib (100 µM) or the respective solvent (DMSO). The y-axis represents colony-forming units per 1 mL of bacterial culture. Results are the mean ± S.D. of four biological replicates (n = 4); *t*-test showed no significant changes between treated and untreated cultures. **E)** Quantification of eDNA in biofilm treated for 24 h with GW4869 (17.3 µM), darapladib (100 µM), and rilapladib (100 µM) or the respective solvent (DMSO) determined with SYBR Green I dye. Results are the mean ± S.D. of four biological replicates (n = 4); *t*-test showed no significant changes between treated and untreated cultures. **F)** Representative confocal scanning microscopy figures of *P. aeruginosa* PA01 grown as static biofilms treated for 24 h with GW4869 (7.5 µM), darapladib (100 µM), rilapladib (100 µM) or DMSO as the solvent control. Biofilms were stained to visualize live (green) or dead/dying (red) cells. **G)** Ratio of dead to live cells in biofilms was determined from red and green fluorescence intensity quantification from Figure 2F using the BiofilmQ tool.

Biofilm formation represents a dominant lifestyle of *P. aeruginosa* during chronic infections (Thi, Wibowo et al. 2020; Tuon, Dantas et al. 2022), functioning as a protective barrier against antibiotics (Soares, Caron et al. 2019; Yin, Cheng et al. 2022). Therefore, we next assessed the effects of GW4869, darapladib, and rilapladib on biofilm formation and cell viability in static cultures (without agitation) adhered to plastic MTP. Pharmaceuticals were introduced at the time of inoculation, followed by a 24-hour incubation at 37 °C under static conditions. Crystal violet staining of adherent cells revealed that these pharmaceuticals had no significant effect on biofilm quantity (Figure 2C). Viable bacterial counts assessed by colony-forming units (CFUs) assays indicated no significant differences between treated and untreated biofilms (Figure 2D). Furthermore, quantification of extracellular DNA, a critical biofilm structural component linked to increased antibiotic resistance (Dai, Luo et al. 2024), showed no significant differences between the treated and untreated biofilms (Figure 2E). Given that the tested pharmaceuticals did not affect initial biofilm formation steps, we next investigated their impact on biofilm dispersal. Using confocal laser scanning microscopy (CLSM), we examined biofilm architecture and cell viability. In this experiment, *P. aeruginosa* biofilms were pre-formed by incubating bacteria at 37 °C under static conditions on microscopic glass slides for 24 h, followed by an additional 24 h treatment with GW4869, darapladib, or rilapladib. Biofilms were then stained with live/dead viability dyes: a green dye for live cells and a red dye for cells with compromised membranes assigned as dead or dying cells. Results showed no discernible effect of the tested pharmaceuticals on biofilm dispersal (Figure 2F). However, the ratio of dead/dying cells was much higher in biofilms treated with all three compounds compared to untreated controls (Figure 2G). Quantitative analysis of CLSM images revealed that the dead-to-live cell ratio increased from 2:1 in untreated biofilms to approximately 3:1 in biofilms treated with darapladib or rilapladib. Notably, GW4869 had a strong effect, as only about 3 % of the cells were estimated to be alive.

These findings highlight the potential of GW4869, darapladib, and rilapladib as inhibitors of planktonic growth and the viability of biofilm-forming *P. aeruginosa* cultures.

### 3.3 Inhibitory potential of selected compounds against PlaF

Next, we investigated whether PlaF, an intracellular PLA recently discovered by our group, serves as a potential target of GW4869, darapladib, and rilapladib. PlaF plays a critical role in maintaining phospholipid homeostasis and impacts the pathogenicity of *P. aeruginosa* (Bleffert, Granzin et al. 2022, Caliskan, Poschmann et al. 2023). To address this, we first performed biochemical assays using purified PlaF reconstituted into phospholipid liposomes to ensure a near-native conformation (Ahmad, Strunk et al. 2021). To assess the inhibition of its hydrolytic activity in the presence of the three selected pharmaceuticals, a spectrophotometric assay based on hydrolysis of the pro-chromogenic *p*-nitrophenyl butyrate (*p*-NPB) substrate revealed that all three compounds inhibited PlaF activity. Darapladib and rilapladib at 400 µM reduced PlaF activity by ∼50 %, while GW4869 (69.2 µM) inhibited PlaF activity by ∼ 30 % (Figure 3A). Although some of the remaining compounds showed an inhibitory effect on the PlaF activity, we did not consider them further as they did not impact growth of *P. aeruginosa*.

**Figure 3.**
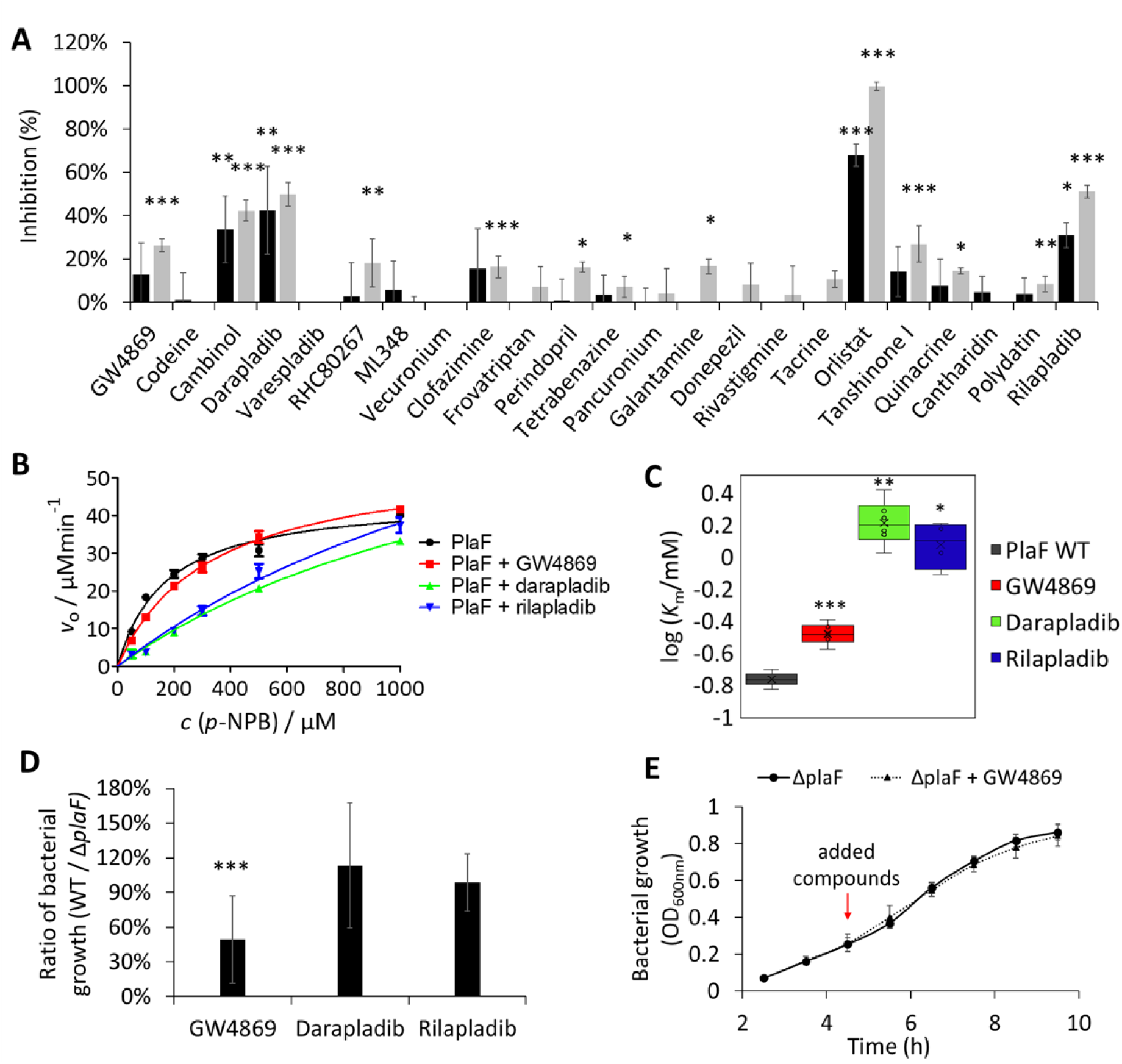
Inhibitory effect of potential PLA targeting pharmaceuticals on PlaF. **A)** Purified PlaF was treated with compounds (34.6 µM and 69.2 µM for GW4869; 66.8 µM and 133.6 µM for codeine; 200 µM and 400 µM for remaining compounds) in the presence of the *p*-NPB substrate, and the released product was quantified spectrophotometrically during 1 h. Black and grey bars indicate lower and higher concentrations of pharmaceuticals, respectively. Results are shown as relative inhibition compared to samples treated with the respective solvent. Results are the mean ± S.D. of three experiments each measured with three replicates (n = 9). *t*-test was calculated by comparing pharmaceutical- and solvent-treated samples, \**p* < 0.05, \*\**p* < 0.01, \*\*\**p* < 0.001. **B)** Michaelis-Menten curve showing enzyme kinetic studies with purified PlaF treated with GW4869 (34.6 µM), darapladib (200 µM), or rilapladib (200 µM) measured spectrophotometrically using *p*-NPB assay. The curves are calculated with GraphPad software from two independent experiments each measured three times (n = 6). **C)** Log-transformed Michaelis-Menten constants calculated from curves in panel B. The average value is indicated with a cross, while the center line represents the median. The box limits indicate the interquartile range, and the whiskers indicate minimum and maximum values. *t*-test of normally distributed values is calculated compared to the non-treated sample, \**p* < 0.05, \*\**p* < 0.01, \*\*\**p* < 0.001. **D)** The ratio of *P. aeruginosa* WT/Δ*plaF* bacterial growth after 6 h incubation with GW4869 (17.3 µM), darapladib (100 µM) and rilapladib (100 µM). Bacterial cultures were grown in LB medium at 37 °C with shaking at 1000 rpm. Results are shown as the ratio of the mean values ± S.D. of two independent experiments each measured three times (n = 6). *t*-test of normally distributed values, \*\*\**p* < 0.001. **E)** Growth curves of PA01 Δ*plaF* treated with GW4869 (17.3 µM). Bacterial cultures were grown in LB medium at 37 °C with shaking at 1000 rpm in plastic MTP. Red arrows indicate time points of compound addition. Results are the mean ± S.D. of four biological replicates (n = 4).

Next, we employed enzyme kinetic studies to determine whether these compounds inhibit PlaF competitively. Using the *p*-NPB assay, we measured PlaF activity at various substrate concentrations (0.05–1 mM) and calculated the initial velocity (*v*₀) and Michaelis-Menten constant (*K*_m_). Kinetic plots showed that darapladib and rilapladib strongly, and GW4869 less prominently affect the catalytic properties of PlaF (Figure 3B) without altering *v*_max_ but increasing the *K*_m_ (Figure 3C). These results indicate that PlaF treated with these compounds exhibits reduced substrate affinity thus suggesting competitive inhibition.

To further explore whether these pharmaceuticals target PlaF in bacteria, we compared the growth of *P. aeruginosa* wild-type and an isogenic Δ*plaF* mutant strain in the presence of the compounds (Figure 3D). Results revealed that darapladib and rilapladib showed no differential effects on the growth of the Δ*plaF* strains, indicating that their mode of action is likely independent of PlaF. In contrast, after 6 h of incubation with GW4869, the Δ*plaF* mutant grew to higher cell density than the wild-type strain (Figure 3D), whereas growth curves of GW4869-treated and -untreated Δ*plaF* did not differ (Figure 3E). These results suggest that the action of GW4869 may be directly or indirectly related to PlaF.

Overall, our findings suggest that GW4869, darapladib, and rilapladib modulate PlaF activity *in vitro*, although a distinct inhibitory effect potentially mediated by inhibition of PlaF in bacterial culture was only observed with GW4869.

### 3.4 Mechanism of inhibition of PlaF by GW4869 involves binding to the substrate-binding tunnel

To obtain atomistic insights into the inhibition of PlaF by GW4869, we conducted unbiased molecular dynamics (MD) simulations. Results from five independent 1 μs long MD simulations indicated that the t-PlaF:GW4869 complex, where the ligand was initially docked at the active site, remains structurally invariant, as indicated by a low overall root mean square deviation (RMSD) of < 1.5 Å (Figure S2). Hierarchical clustering of the simulated configurations sampled every 50 ps across the five replicas revealed that GW4869 preferentially binds to tunnels T2 and T3, which are structural elements previously identified as essential for substrate hydrolysis (Ahmad, Strunk et al., 2021; Gentile, Modric et al., 2024). The most abundant cluster (C1), where GW4869 occupies parts of T2 and T3, accounted for 25.7 ± 4.2% of the analyzed frames (Figure 4A).

**Figure 4:**
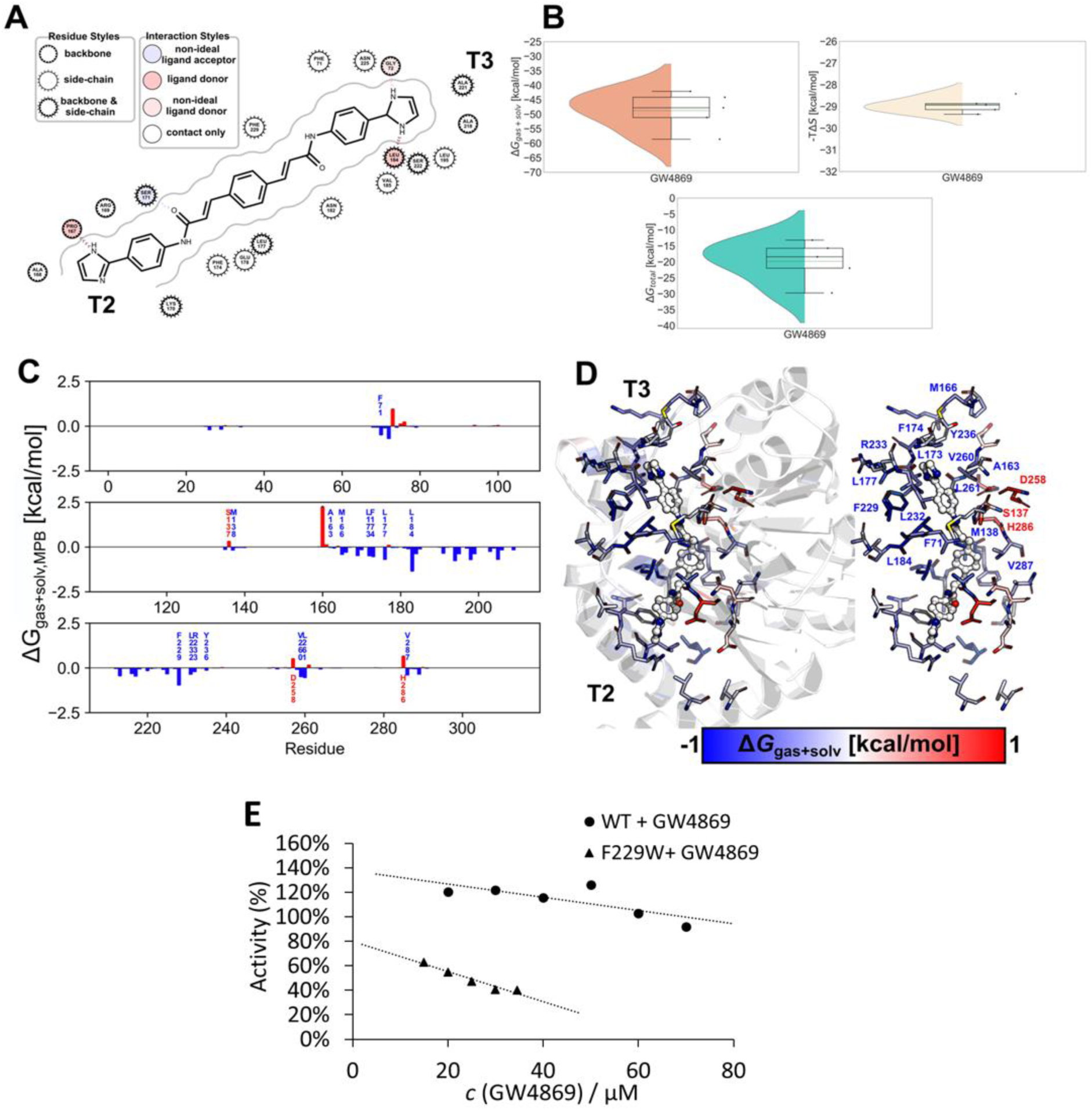
Molecular mechanism of GW4869 binding to t-PlaF. **A)** Binding of GW4869 to t-PlaF in the most populated cluster from MD simulations represented as 2D-interaction map. Residues interact with their side-chain, backbone or both, as detailed by the legend. The entrances of tunnels T2 and T3 are indicated. **B)** Effective energy (top left) and configurational entropy (top right) contributions to binding, and total binding energy (bottom) of GW4869. The violin plots indicate the distributions of the data points; the inner box of a box plot represents the interquartile range, with the horizontal black line indicates the median. The dotted green line represents the mean, and the whiskers show the rest of the distribution, excluding points determined to be outliers when they fall outside 1.5 times the interquartile range. **C)** Per-residue binding effective energy of tunnel residues. The energies were averaged across five independent replicas (n = 5). Error bars denote the SEM. **D)** Per-residue binding effective energy mapped at the structural level. Interacting residues are depicted as sticks, colored according to the effective binding energy. The entrances of tunnels T2 and T3 are indicated. The interacting residues of T3 are labeled. The GW4869 inhibitor is depicted as red, blue, and grey spheres. **E)** Determination of half-maximal inhibitory concentration for inhibition of phospholipid liposome-reconstituted PlaF_WT_ or PlaF_F229W_ with GW4869 using *p*-NPB assay. Graphs show linear plots of relative activities compared to the untreated sample set to 100 %. The result represents the mean ± S.D. of six measurements (n = 6).

To quantify the binding energy of GW4869 in cluster C1, the unbiased MD simulations were used for molecular mechanics Poisson-Boltzmann surface area (MM-PBSA) calculations, incorporating an implicit membrane model (Miller, McGee et al., 2012; Greene, Qi et al., 2019). The computations were converged, as evidenced by the comparison between the first and the second halves of the trajectories (Figure S3). These trajectories yielded a favorable binding effective energy (Δ*G*_gas+solv_) of -48.8 ± 2.9 kcal mol^−1^ and an unfavorable entropic contribution to binding (*T*Δ*S*) of -28.9 ± 0.2 kcal mol^−1^. The resulting binding free energy (Δ*G*_bind_) estimate is -19.8 ± 2.9 kcal mol^−1^ (Figure 4B). The corrected standard free energy of binding (Δ*G*^0^) is -21.7 ± 2.9 kcal mol^−1^ indicating that binding of GW4869 to T2 and T3 is exergonic.

The residue-specific decomposition of the binding effective energy revealed key residues in T2 and T3 that contribute significantly to GW4869 binding. Residues in T3 that were proposed to be part of the product egress pathway (Ahmad, Strunk et al. 2021) contributed more to GW4869 binding (62.2 % of the total favorable interacting residues) than residues in T2 (37.8 %). This resulted in a more favorable interaction energy with T3 (Δ*G*_gas+solv, T3_ = -30.4 ± 2.2 kcal mol^−1^) than T2 (Δ*G*_gas+solv, T2_ = -15.0 ± 3.2 kcal mol^−1^). Notably, residue F229 in T3, previously shown to be critical for product release, exhibited one of the strongest binding interactions. Other residues near F229, including F71, L177, L184, and L232, also contributed markedly to GW4869 binding (Figure 4C-D).

To experimentally validate the role of F229 in GW4869 binding, we aimed to determine the half-maximal inhibitory concentration (IC_50_) of GW4869 for wild-type PlaF (PlaF_WT_) and the F229W variant (PlaF_F229W_). Both variants were purified and reconstituted into phospholipid liposomes to ensure near-native PlaF conformations (Ahmad, Strunk et al. 2021). Inhibition curves obtained by varying the GW4869 concentration at constant protein and substrate (*p*-NPB) concentrations displayed a linear relationship within the tested concentration range without reaching a plateau, likely due to the low solubility of the hydrophobic GW4869 in water and its adsorption to phospholipid liposomes. Although precise IC_50_ values could not be determined, the observed linear relationship suggests that GW4869 inhibits PlaF_F229W_ activity more efficiently compared to PlaF_WT_ within the tested concentration range (Figure 4E). We speculate that the increased indolyl π-system of Trp and/or the presence of a hydrogen bond donor in the Trp ring provide more favorable interactions with GW4869 compared to the smaller phenyl ring of phenylalanine.

In summary, these findings suggest that GW4869 binds noncovalently to the active site of PlaF, which might lead to interference with substrate binding and product release.

### 3.5. Compounds inhibiting *P. aeruginosa* growth potentiate the effect of common antibiotics

To evaluate potential synergistic effects of *P. aeruginosa* growth-inhibiting pharmaceuticals GW4869, darapladib, and rilapladib (Figure S4) with common antibiotics against *P. aeruginosa*, we conducted combinatorial treatments with antibiotics of differing modes of action (Yayan, Ghebremedhin et al. 2015). The selected last-resort antibiotics included gentamicin, which targets ribosomal function (Tangy, Moukkadem et al. 1985); colistin, which disrupts cytoplasmic membrane integrity (Falagas, Kasiakou et al. 2005); and piperacillin and imipenem, which target cell wall synthesis (Lipman and Neu 1988, Perry and Markham 1999). To assess synergistic effects, these bactericidal antibiotics were used at sub-inhibitory concentrations for *P. aeruginosa*. The concentrations used were as follows: gentamicin at 0.5 mg/L (0.70 µM), piperacillin at 2 mg/L (3.71 µM), colistin at 1 mg/L (0.71 µM), and imipenem at 2 mg/L (6.30 µM) (Yayan, Ghebremedhin et al. 2015).

Treatment of planktonic *P. aeruginosa* PA01 cultures in the exponential growth phase with combinations of pharmaceuticals and antibiotics revealed that darapladib did not enhance the activity of any of the four antibiotics tested (Figure S5). Similarly, GW4869 and rilapladib did not improve the bactericidal effects of gentamicin, colistin, or piperacillin (Figure S5). However, combinations of GW4869 or rilapladib with imipenem showed significant synergy (Figure 5A). Imipenem at a concentration four-fold lower than its minimum inhibitory concentration had only minimal impact on bacterial growth. In contrast, co-treatment with GW4869 (17.3 µM) or rilapladib (100 µM) abolished *P. aeruginosa* growth within two hours and significantly decreased the optical density after an additional three hours (Figure 5A).

**Figure 5.**
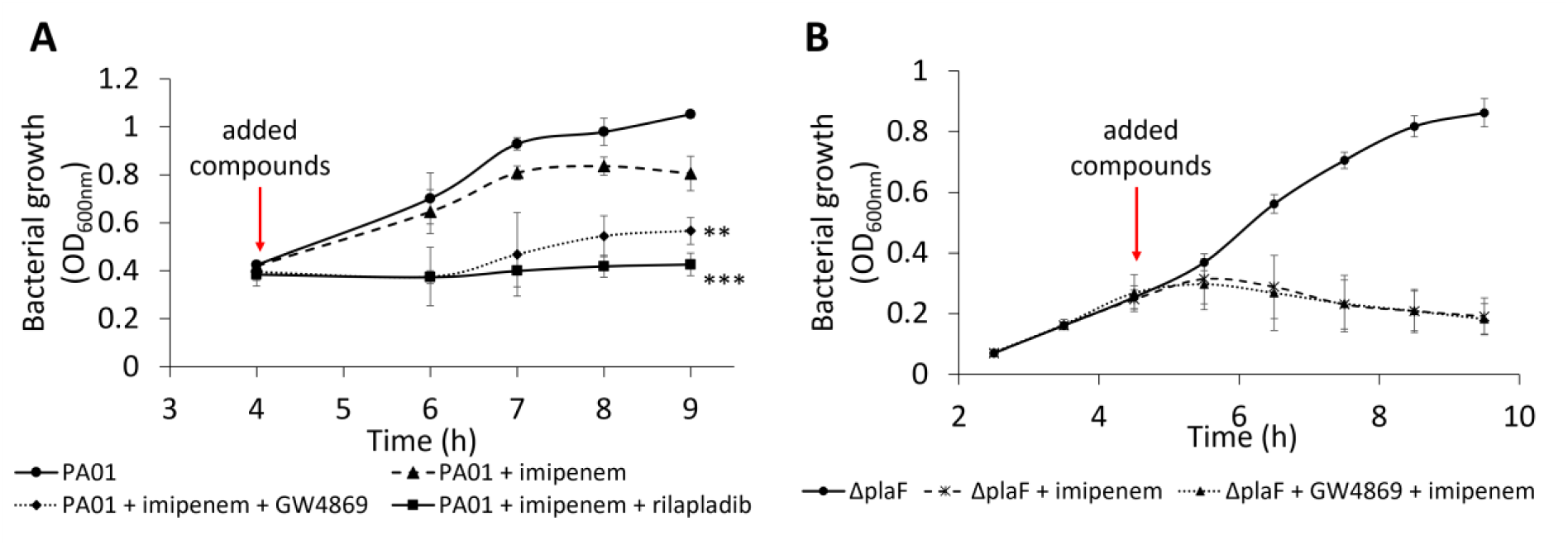
PlaF inhibitors potentiate the antibiotic activity of imipenem on *P. aeruginosa*. **A)** The combination of imipenem (6.3 µM) and GW4869 (17.3 µM) or rilapladib (100 µM) resulted in reduced growth of *P. aeruginosa.* Bacterial cultures were grown in LB medium at 37 °C with shaking at 1000 rpm. Results are shown as the mean ± S.D. of four biological replicates (n = 4). *t*-test of normally distributed values, ** *p* < 0.01, *** *p* < 0.001. **B)** Growth of *P. aeruginosa* Δ*plaF* mutant in the presence of a combination of imipenem (6.3 µM) and GW4869 (17.3 µM). Bacterial cultures were grown in LB medium at 37 °C with shaking at 1000 rpm. Results are shown as the mean ± S.D. of four biological replicates (n = 4).

To explore the mechanism of action of GW4869, presumed to inhibit PlaF, we examined its effect in combination with imipenem on the Δ*plaF* mutant strain (Figure 5B). Growth analyses indicated that the GW4869-imipenem combination had no different effect than imipenem alone. In both cases, growth was slowed within one hour of treatment, with no subsequent growth over the next four hours. These findings suggest that GW4869’s synergism with imipenem is dependent on the presence of PlaF, further supporting the hypothesis that GW4869 act on *P. aeruginosa* by inhibiting PlaF.

## 4. Discussion

### Inhibition of intracellular phospholipase A is a novel strategy to combat Pseudomonas aeruginosa virulence

Various virulence factors of *P. aeruginosa* play a crucial role in host infections by facilitating bacterial adhesion, invasion, immune suppression, tissue damage, and nutrient acquisition (Liao, Huang et al. 2022). These mechanisms highlight the potential of antivirulence strategies as innovative therapeutic approaches (Veesenmeyer, Hauser et al. 2009). This study investigates whether inhibiting intracellular phospholipase PlaF, a virulence factor of *P. aeruginosa*, can suppress growth of bacterial culture *in vitro*, providing a foundation for developing antivirulence therapies targeting intracellular PLAs.

Intracellular PLAs are essential for remodeling the bacterial membrane phospholipid composition, a vital adaptation that enhances virulence and antibiotic resistance in *P. aeruginosa* (Bleffert, Granzin et al. 2022, Caliskan, Poschmann et al. 2023) and other pathogens (Kerrinnes, Young et al. 2015). Using a target-based drug repurposing strategy, 23 potential PLA inhibitors were selected from compounds already approved or undergoing (pre-)clinical trials for human diseases. This approach could leverage the safety profiles of these compounds, including data on toxicity, pharmacokinetics, and reproductive and carcinogenic effects (Mishra, Vasanthan et al. 2024). While cross-reactivity with human pathways could pose challenges, it may also offer dual benefits by targeting pathogen factors and host pathways critical for infection resolution or damage mitigation (Hassan and Blanchard 2022). These inhibitors could also serve as leads for developing bacteria-specific drugs targeting intracellular PLAs.

### Efficacy of PLA inhibitors in inhibiting P. aeruginosa growth

Our results reveal that GW4869, darapladib, and rilapladib significantly inhibit the *in vitro* growth of planktonic *P. aeruginosa* and decrease the viability of biofilm-forming cells. While for rilapladib and darapladib, targeting human lipoprotein-associated phospholipase A2 (Corson 2010, Maher-Edwards, De’Ath et al. 2015), no clearly defined bacterial target could be identified, GW4869 demonstrated potent inhibition of the intracellular PLA PlaF. This was supported by the resistance observed in *P. aeruginosa ΔplaF* mutants and the inhibition of purified PlaF. GW4869 reduced planktonic *P. aeruginosa* growth more than 50% within seven hours and drastically reduced biofilm-cell viability to 3% post-treatment. These findings align with prior studies showing impaired biofilm formation in *plaF* knockout strains (Bleffert, Granzin et al. 2022). Interestingly, GW4869 had no significant impact on *Legionella pneumophila* replication or outer membrane vesicle (OMV) secretion in infected THP-1 cells (Jung, Herkt et al. 2017), consistent with the absence of PlaF homologs in *L. pneumophila* (Flores-Díaz, Monturiol-Gross et al. 2016).

#### Synergy with antibiotics

GW4869 also demonstrated notable adjuvant activity when combined with imipenem, even at sub-minimum inhibitory concentrations (MIC). The synergy might result from reduced production of the imipenem neutralizing β-lactamase AmpC (Lister, Wolter et al. 2009) in the Δ*plaF* mutant (Bleffert, Granzin et al. 2022). Imipenem disrupts peptidoglycan synthesis by inhibiting penicillin-binding proteins, destabilizing the cell wall (Lister, Wolter et al. 2009). Thus, a dual assault on the bacterial envelope due to a putative membrane dysfunction caused by GW4869-mediated PlaF inhibition might enhance bactericidal activity.

### Potential alteration of host-pathogen interactions

Rilapladib and darapladib inhibit host lipoprotein-associated PLA2 which release precursors of inflammatory lipid mediators (Maher-Edwards, De’Ath et al. 2015; Corson 2010, Zhang, Xu et al. 2016, Zhuo and Yuan 2020) by rilapladib or darapladib would help dampening systemic inflammation and simultaneously slower bacterial proliferation thus helping clearing infections.

GW4869 is a noncompetitive inhibitor of neutral sphingomyelinase 2 (nSMase2), which regulates the conversion of sphingomyelin to ceramide (Luberto, Hassler et al. 2002; Tabatadze, Savonenko et al. 2010; Wan, Fang et al. 2023). This mechanism has been shown to inhibit exosome release, which results in impaired pro-tumorigenic macrophage differentiation (Peng, Zhao et al. 2022) and suppression of systemic inflammation in murine sepsis model and LPS treatment (Peng, Zhao et al. 2022). These immunomodulatory effects, coupled with its direct activity against *P. aeruginosa*, suggest that GW4869 may bolster host defenses against severe bacterial infections.

Another intriguing mode of GW4869 action may involve the manipulation of ceramide, which was shown to promote infection and immune evasion (Grassmé and Becker 2013; Duarte, Akkaoui et al. 2020). Bacteria do not synthesize sphingomyelin or ceramide, but they release ceramide through enzymatic hydrolysis of host sphingomyelin by sphingomyelinases, e.g., *Staphylococcus aureus* β-hemolysin and *Clostridium perfringens* α-toxin (Flores-Díaz, Monturiol-Gross et al. 2016). While *P. aeruginosa* lacks sphingomyelinase, it produces phospholipase C, a virulence factor capable of hydrolyzing sphingomyelin *in vitro*. It remains to be seen whether GW4869 mediates ceramide reduction in *P. aeruginosa* and whether this could interfere with host-pathogen interactions that promote infection and immune evasion.

### Future directions

Further preclinical and clinical research is required to determine whether GW4869 could be effectively used alongside antibiotics to improve outcomes in bacterial infections. GW4869 holds promise as an antivirulence compound targeting intracellular bacterial PLAs and host lipid pathways critical to host-pathogen interactions. These findings open novel avenues for the development of innovative therapies targeting intracellular PLAs to combat bacterial infections.

## 6. Acknowledgments

H.G. is grateful to OpenEye for a Free Public Domain Research License. We are grateful for computational support and infrastructure provided by the “Zentrum für Informations-und Medientechnologie” (ZIM) at the Heinrich Heine University Düsseldorf. We gratefully acknowledge the Gauss Centre for Supercomputing e.V. (www.gauss-centre.eu) for funding this project (user ID: t1ss) by providing computing time on the GCS Supercomputer JUWELS (Alvarez 2021) at Jülich Supercomputing Centre (JSC).

